# Intrauterine growth restriction is associated with adaptive hematopoietic reprogramming and selective immune rewiring in monozygotic twins

**DOI:** 10.64898/2026.08.03.742402

**Authors:** Hyeji Kang, Soonjoung Kim, Sohee Kim, Ji Hoi Kim, Chan-Wook Park, Joong Shin Park, June-Yong Lee, Dongsung Lee, Jong Kwan Jun, Seung Mi Lee, Chul-Hwan Lee

## Abstract

**Background:** Prenatal growth restriction has been associated with adverse neonatal and long-term health outcomes, yet the epigenetic mechanisms by which an adverse intrauterine environment shapes fetal immune development remain incompletely understood. Monozygotic dichorionic-diamniotic twins with selective fetal growth restriction (sFGR) provide a unique human model for investigating environmentally driven developmental programming independent of genetic variation and inter-twin placental vascular anastomoses.

**Methods:** Umbilical cord blood buffy coat samples were collected from three sFGR and two gestational age-matched concordant control twin pairs. Bulk RNA sequencing and genome-wide DNA methylation analysis were performed, followed by differential expression, pathway enrichment, hematopoietic and immune module analyses, differential methylation, and integrative transcriptomic-epigenomic analyses.

**Results:** Compared with concordant control twin pairs, discordant twins exhibited broad attenuation of immune and inflammatory transcriptional programs alongside enrichment of erythroid- and hypoxia-related pathways, consistent with adaptive hematopoietic responses to intrauterine stress. Within discordant twin pairs, the growth-restricted co-twins displayed marked transcriptional asymmetry characterized by selective enrichment of cytotoxic lymphoid signatures despite global suppression of myeloid and antigen-presenting cell-associated programs. Integrated transcriptomic and epigenomic analyses further revealed coordinated epigenetic remodeling, with hypomethylated regions in growth-restricted twins enriched for immune regulatory pathways, including T cell differentiation and leukocyte activation. At selected loci, concordant hypomethylation and increased gene expression suggested a potential epigenetic basis for the observed immune remodeling.

**Conclusions:** These findings suggest that intrauterine growth restriction is associated with coordinated hematopoietic and immune reprogramming at birth, consistent with both compositional and cell-intrinsic alterations. In genetically identical twins, relative growth divergence was associated with polarized transcriptional states, highlighting how intrauterine environmental differences may shape early immune development independent of genetic background.

## Backgrounds

Disentangling environmental factors from genetic influences remains a central challenge in human studies of complex traits and diseases. Monozygotic (MZ) twin studies provide a powerful natural framework for addressing this challenge, as co-twins share an essentially identical genome, enabling phenotypic differences to be attributed primarily to non-genetic factors [1–4]. However, MZ twins do not always experience identical prenatal environments. The timing of embryonic splitting determines placental and amniotic architecture, resulting in distinct chorionicity types—dichorionic-diamniotic (DCDA), monochorionic-diamniotic, and monochorionic-monoamniotic—that impose different degrees of intrauterine heterogeneity [5–11]. In DCDA twins, early embryonic splitting leads to the formation of independent placentas and amniotic sacs, exposing each fetus to a distinct local intrauterine environment despite their shared genetic identity [12, 13]. This naturally occurring prenatal divergence provides a unique opportunity to examine how distinct intrauterine environments shape fetal immune and hematopoietic transcriptional programs.

One of the most readily observable manifestations of this variability is birthweight discordance. Although fetal growth is strongly influenced by genetic factors [14], substantial birthweight differences between monozygotic twins are often linked to intrauterine conditions rather than DNA sequence variation. In particular, placental insufficiency, unequal placental development [15], maternal nutritional status [16], and localized differences in uteroplacental perfusion have all been implicated as key contributors. Birthweight discordance is associated with disparities in neonatal survival [17], endocrine profiles [18], immune maturation, and long-term neurodevelopmental outcomes [19]. Notably, however, this discordance is typically not inherited from the parents [20]. The absence of heritability suggests an acquired molecular basis, positioning epigenetic and transcriptional regulation as central mediators linking the prenatal environment to postnatal phenotypic variation.

Among epigenetic mechanisms, DNA methylation is particularly sensitive to intrauterine nutritional deprivation during fetal development, and such marks can persist as epigenetic memory throughout life [21, 22]. DNA methylation has also been implicated in the long-term regulation of growth, metabolic, and immune cell function [22, 23], supporting its potential role in fetal immune programming. However, whether intrauterine growth divergence in genetically identical human twins induces coordinated immune and hematopoietic reprogramming remains unclear.

In monochorionic twin pregnancies, environmentally driven phenotypic divergence, such as sFGR [24, 25] and twin-to-twin transfusion syndrome [26], have been extensively studied as consequences of shared placental vasculature and imbalanced blood flow [27]. In contrast, growth discordance in MZ-DCDA twins arises in the absence of shared placental circulation and instead reflects localized placental function and implantation-related factors. This distinction makes MZ-DCDA twins a particularly powerful system for capturing the transcriptional consequences of prenatal growth divergence without the confounding effects of direct vascular anastomoses.

Despite the recognized importance of early-life programming, most prior studies using disease-discordant monozygotic twins have focused on postnatal or adult phenotypes and have often relied on targeted epigenetic analyses [28, 29]. Genome-wide investigations of transcriptional and epigenetic states at birth, when prenatal environmental effects are most directly captured, remain remarkably limited [30]. In particular, integrative analyses of gene expression and DNA methylation in umbilical cord blood–derived immune cells in monozygotic twins discordant for birthweight are scarce [31], leaving a critical gap in our understanding of how prenatal growth differences are molecularly encoded during early immune development.

In this study, we leverage MZ-DCDA twin pregnancies as a unique human model to define neonatal transcriptional signatures associated with prenatal growth discordance. By performing RNA sequencing (RNA-seq) on the buffy coat isolated from umbilical cord blood at birth, we directly captured fetal transcriptional programs shaped by distinct growth trajectories within genetically identical pairs. Through comparison between concordant twins and twins with sFGR, alongside intra-pair analyses of normal and growth-restricted co-twins, we sought to identify transcriptional programs reflecting immune and hematopoietic adaptations associated with unequal prenatal growth. In addition, we performed Whole-Genome Bisulfite Sequencing (WGBS) to analyze DNA methylation differences between discordant co-twins and to investigate whether selective immune remodeling in growth-restricted twins is accompanied by corresponding epigenetic alterations. This integrative approach enables us to delineate transcriptional and epigenetic signatures associated with early growth differences, providing insight into the molecular mechanisms by which prenatal microenvironmental variation influences neonatal immune development.

## Methods

### Subjects, sample collection, and clinical assessment

The case group consisted of six dichorionic monozygotic twin neonates (three twin pairs) with sFGR, defined as small-for-gestational-age birthweight in one twin and normal birthweight in the co-twin. The control group consisted of four dichorionic monozygotic twin neonates (two twin pairs) with concordant birthweight, matched for maternal age and gestational age at delivery. Chorionicity was determined by clinical ultrasound in early pregnancy, and zygosity was confirmed by short tandem repeat (STR) analysis using cord blood from each twin fetus, as previously described [32]. Umbilical cord blood was collected into EDTA tubes immediately after delivery, and buffy coat was isolated by centrifugation and stored at −80°C for subsequent RNA and DNA extraction.

### Total RNA extraction and library preparation

Total RNA was extracted using the TRIzol reagent (Thermo Fisher Scientific, 15596018). Globin transcripts were depleted using the TruSeq Stranded Total RNA with Ribo-Zero Globin kit (Illumina, RS-122-2501), and sequencing libraries were constructed using the SureSelectXT RNA Direct Library Preparation kit (Agilent Technologies, G7564), a capture-based, ORF- enrichment approach selected to ensure high-quality data from low-RIN samples. Libraries were sequenced on an Illumina platform in 101 bp paired-end mode.

### RNA-seq processing and differential expression analysis

Raw reads were trimmed using Trimmomatic (v0.38) and aligned to GRCh38 using HISAT2 (v2.1.0). SAM files were converted to sorted BAM files using SAMtools (v1.22), then to CPM-normalized bigWig tracks using deepTools (v3.5.6) for visualization in IGV (v2.18.4). Gene-level counts were generated using featureCounts (Rsubread) against the Ensembl GRCh38.114 annotation, retaining genes with ≥10 total counts. Differential expression was assessed using DESeq2 (v1.50.2): We first compared twins from discordant pregnancies with gestational age– matched concordant controls using an unpaired analysis to identify transcriptomic features associated with growth-discordant pregnancies. We then performed paired analyses within discordant twin pairs to isolate transcriptional changes specific to the growth-restricted co-twin while controlling for shared genetic background. To identify transcriptional differences associated with growth discordance at the group level, an unpaired design was used to compare all twins from discordant pregnancies with twins from concordant control pregnancies. To identify transcriptional differences associated specifically with sFGR within genetically matched twin pairs, a paired design was used in which twin pair (Parent) was included as a blocking factor (∼ Parent + weight_class). Adjusted p-values were calculated using the Benjamini-Hochberg (BH) method, with significance defined as adjusted p<0.05 and |log2(fold change)|>1.

### Gene set enrichment, pathway, and module score analysis

Pathway-level enrichment analysis was conducted using rank-based approaches applied to the full transcriptome. Genes were ranked by the DESeq2 Wald statistic, and Gene Set Enrichment Analysis (GSEA) was performed using fgsea (v1.36.2) with MsigDB gene sets obtained via msigdbr (v25.1.1), including Hallmark and C2 curated pathways. Enrichment significance was assessed using BH-adjusted p-values. Gene Ontology (GO) enrichment analysis was performed using the gseGO function from clusterProfiler (v4.18.4), focusing on Biological Process (BP) terms; enriched GO terms were ranked by normalized enrichment score (NES), and semantically redundant terms were reduced using the simplify function in clusterProfiler.

Module-based transcriptional profiling was performed to evaluate lineage- and immune-related programs across concordant control, Disc-N, and Disc-sFGR twin samples, Marker gene sets representing major hematopoietic and immune lineages were manually curated from well-established canonical lineage markers commonly used in human hematopoietic and single-cell transcriptomic studies. Each module comprised a small set of lineage-defining genes selected to capture coordinated biological programs rather than individual differentially expressed genes.

Module scores were calculated for each sample by averaging the scaled expression of genes within each module and Z-score transformed across samples for heatmap visualization, with group-wise differences in module activity visualized using box plots for individual lineage programs. To statistically evaluate coordinated enrichment of predefined marker sets, competitive gene set testing was performed using the CAMERA method implemented in the limma package [33]. Genes were ranked according to the moderated *t*-statistic from the corresponding limma linear model, and enrichment of each lineage-specific marker set was visualized using barcode plots.

### Genomic DNA extraction and library preparation

Genomic DNA was extracted using the QIAamp DNA Blood Mini Kit (QIAGEN, 51104). Sequencing libraries were constructed using the Accel-NGS Methyl-Seq DNA Library Kit (Swift Biosciences, 30096) according to its protocol and sequenced on an Illumina platform using 150 bp paired-end sequencing.

### WGBS data processing and differential methylation analysis

Reads were quality-filtered using Trim Galore (v0.6.10) and aligned to the T2T-CHM13v2.0 (hs1) genome using Bismark (v0.24.2)/Bowtie2 (v2.4.4; --score_min L,0,-0.2). PCR duplicates were removed (deduplicate_bismark), and CpG methylation was extracted using Bismark’s methylation extractor.

Differential methylation between Disc-N and Disc-sFGR twins (n=54,227,132 CpG loci, coverage ≥1 in all six samples) was assessed using DSS (v2.56.0) via bsseq (v1.44.1). A paired linear model (∼ parent + weight) accounting for twin-pair identity as a blocking factor was fitted using DMLfit, and differentially methylated loci (DMLs) were identified at each CpG site using a Wald test via DMLtest. Differentially methylated regions (DMRs) were called using callDMR (p.threshold=0.1, minlen=30 bp, minCG=2, dis.merge=150 bp) and filtered by |Δmethylation|≥20%, yielding 2,165 DMRs (1,104 hypermethylated and 1,061 hypomethylated in sFGR). Of these, 2,140 DMRs (1,093 hypermethylated and 1,047 hypomethylated) were retained for downstream analyses after excluding DMRs with missing methylation values in any sample.

### DMR annotation, GO enrichment, and integration with RNA-seq data

DMRs were annotated using ChIPseeker (v1.44.0) against the hs1 NCBI RefSeq transcript database (TSS: −2,000 to +200 bp); for DMRs overlapping intronic or exonic regions, the gene identifier was corrected to the physically overlapping gene rather than the nearest TSS-based assignment.

GO Biological Process enrichment of DMR-annotated genes (vs all human genes) was performed using clusterProfiler (v4.16.0) with org.Hs.eg.db (v3.21.0); significance was assessed using BH-adjusted p-values (p<0.05).

DMR-annotated genes were compared with paired RNA-seq DEGs (Disc-N vs Disc-sFGR; DESeq2, adjusted p<0.05, |log2(fold change)|>1) by gene symbol to identify loci with concordant epigenetic and transcriptional changes.

### Cell-type deconvolution analysis

Cell-type fractions in WGBS samples were estimated using EpiDISH (v2.24.0)[34] with the centUniLIFE.m reference panel[35], a unified DNA methylation reference covering 19 immune cell types applicable to blood tissue from birth to old age, defined by 1,906 CpG probes derived from the Illumina 450K/EPIC methylation array (hg19 genome build). Reference probe coordinates were converted from hg19 to hs1 using the rtracklayer (v1.68.0) and the UCSC hg19-to-CHM13v2 liftOver chain, and all 1,906 probes were uniquely mapped. For each probe, the methylation (M) and total coverage (Cov) values were summed on both strands of the CpG dyad from the WGBS data, and probes with a coverage of 1 or greater in all samples (1,905 out of 1,906) were retained. Using the robust partial correlation (RPC) method implemented in EpiDISH, we first estimated the proportions by cell type using the entire reference data for 19 cell types. Since the proportions of 12 adult-specific cell types were estimated to be negligibly low, we subdivided the reference data into 7 cord blood-related cell types (granulocytes, CD4+ T cells, CD8+ T cells, B cells, monocytes, NK cells, and nucleated red blood cell) and used this subset to re-estimate the proportions for analysis.

### Data visualization

Volcano, GO dot, DMR lollipop, and PCA plots were generated using ggplot2 (v4.0.3)/ggrepel (v0.9.8); lollipop gene tracks were captured from IGV using the T2T-CHM13v2.0 (hs1) assembly and NCBI RefSeq annotation. Heatmaps (pheatmap/ComplexHeatmap v2.24.1) used row-wise Z-score-transformed expression or methylation values, and Venn diagrams were generated using ggvenn (v0.1.19).

### Use of generative artificial intelligence and figure preparation

ChatGPT (OpenAI) was used to generate initial illustrative elements for Figures 1A and 4E. All generated content was reviewed, modified, and approved by the authors. Final figure assembly was performed using BioRender.com.

**Figure 1.**
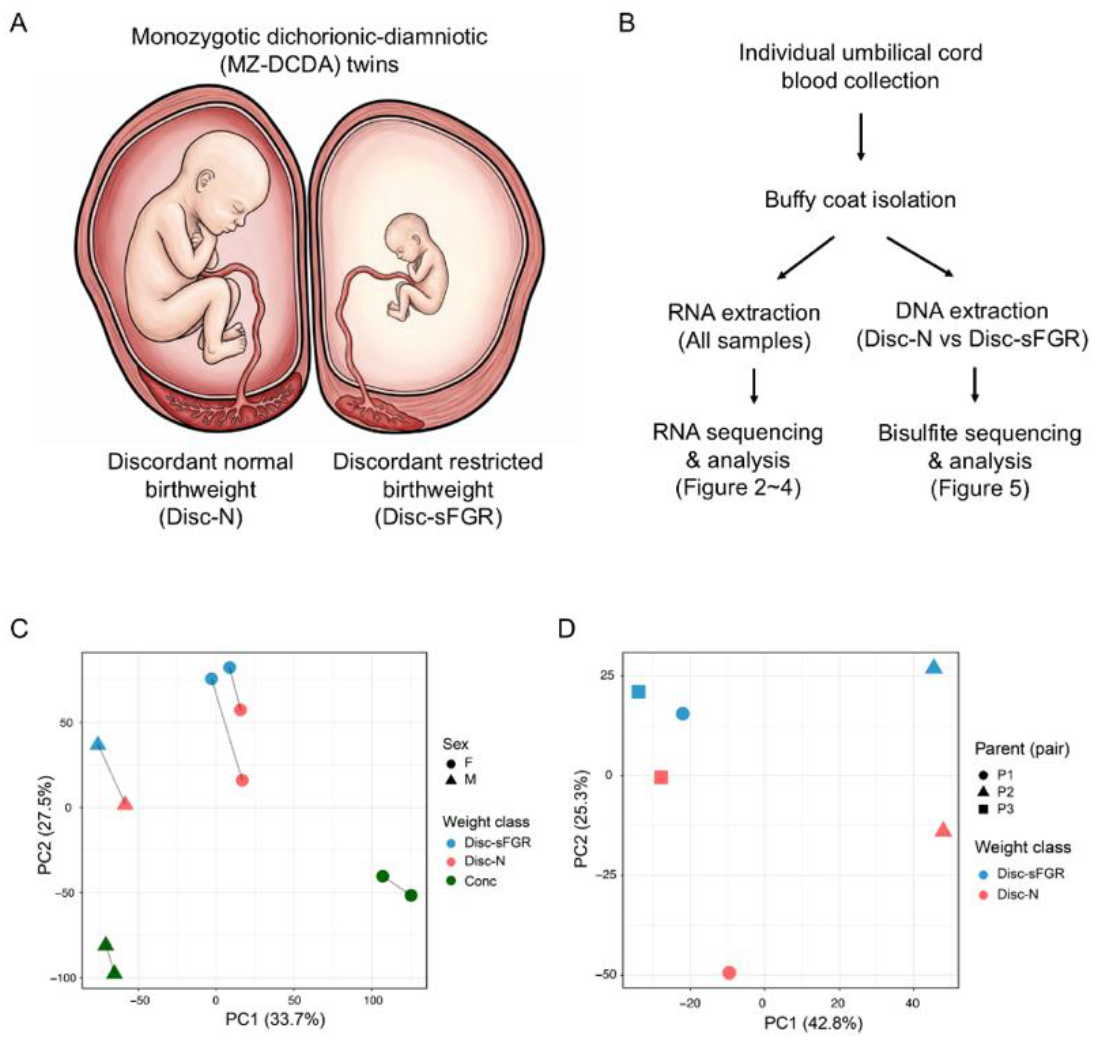
Study design and global transcriptional landscape of MZ-DCDA twin pregnancies with sFGR. (A) Schematic illustration of selective fetal growth restriction (sFGR) in monozygotic dichorionic–diamniotic (MZ-DCDA) twin pregnancies, highlighting the independent placentas that generate distinct intrauterine environments despite identical genetic backgrounds. (B) Overview of the study workflow, including neonatal sample collection, RNA sequencing, Whole-genome bisulfite sequencing and comparative transcriptomic and DNA methylation analyses. (C) Principal component analysis (PCA) of neonatal transcriptomes showing distinct clustering of discordant-sFGR (Disc-sFGR), discordant-normal (Disc-N), and concordant (Conc) samples. (D) PCA of neonatal transcriptomes from Disc-sFGR and Disc-N twins, excluding concordant samples.

## Results

### A genetically controlled twin model reveals global transcriptional alterations associated with fetal growth restriction

We studied a rare group of MZ-DCDA twin pairs enrolled in the Seoul National University Hospital twin cohort (Figure. 1A and 1B) to identify biological differences associated with fetal growth restriction independent of genetic background. The cohort included pregnancies complicated by sFGR as well as concordant control pairs, enabling genetically matched comparisons while minimizing the confounding effects associated with placental sharing. Twin pairs were classified as having selective fetal growth restriction (sFGR) if the intertwin birthweight discordance exceeded 20% [36]. Cases where there are no significant intertwin growth discordance and their gestational ages are similar were designated as concordant controls (Conc). Importantly, gestational age at delivery was comparable between concordant and discordant groups, allowing growth-related transcriptional differences to be examined independently of gestational age. Within sFGR pairs, co-twins were further categorized by birthweight status as the discordant-normal (Disc-N) twin and the discordant-sFGR (Disc-sFGR) twin. This terminology was used consistently throughout the study, and the clinical characteristics of the study cohort are summarized (Table 1).

**Table 1.** Clinical characteristics and sequencing information of the study cohort, including individual birthweights of sFGR and concordant control twin pairs.

| Sample ID | Study group | Twin pair | Maternal age, years | Gestational age at delivery, (weeks/days) | Birth weight, g | Sex | Discordance, % | RNA-seq | WGBS |
| --- | --- | --- | --- | --- | --- | --- | --- | --- | --- |
| 1-sFGR | Discordant | Parents 1 | 34 | 37 weeks/ 3 days | 2100 | F | 23.1 | Yes | Yes |
| 1-N |  |  |  |  | 2730 | F |  | Yes | Yes |
| 2-sFGR |  | Parents 2 | 38 | 37 weeks/ 2 days | 1960 | M | 24.5 | Yes | Yes |
| 2-N |  |  |  |  | 2595 | M |  | Yes | Yes |
| 3-sFGR |  | Parents 3 | 32 | 36 weeks/ 6 days | 1910 | F | 20.7 | Yes | Yes |
| 3-N |  |  |  |  | 2410 | F |  | Yes | Yes |
| 4-C1 | Concordant | Parents 4 | 34 | 37 weeks/ 2 days | 2660 | F | 0.0 | Yes | No |
| 4-C2 |  |  |  |  | 2660 | F |  | Yes | No |
| 5-C1 |  | Parents 5 | 34 | 37 weeks | 2510 | M | 6.0 | Yes | No |
| 5-C2 |  |  |  |  | 2670 | M |  | Yes | No |

We performed bulk RNA sequencing on the buffy coat isolated from umbilical cord blood at birth, rather than Peripheral Blood Mononuclear Cell (PBMC), to preserve granulocytes along with mononuclear leukocytes and platelets and thereby capture a more comprehensive representation of circulating immune cells. Across all samples, quality control metrics indicated robust and highly comparable transcriptome profiles despite the use of frozen buffy coat–derived material (Figure. S1A-S1D). Sample-to-sample correlations were uniformly high, and global expression distributions showed minimal variability, with nearly identical VST profiles across all samples. These results support the conclusion that RNA integrity and sequencing quality were well preserved, enabling reliable downstream comparative analyses. Principal component analysis of transcriptomic profiles demonstrated distinct clustering between concordant and discordant groups, indicating that both twins within the discordant group exhibit globally distinct transcriptomic signatures compared with concordant controls (Figure. 1C). Within the discordant pairs, further separation was observed between Disc-N and Disc-sFGR co-twins, highlighting transcriptional divergence associated with intra-pair growth discordance (Figure. 1D). In contrast, concordant twin pairs clustered tightly together, reflecting highly similar transcriptomic profiles and minimal intra-pair variation (Figure. 1C). Notably, the birthweights of Disc-N twins were comparable to those of concordant twins, suggesting that the observed transcriptional differences are not attributable solely to absolute birthweight. Rather, these findings support the notion that differential intrauterine environments associated with sFGR shape systemic immune-related transcriptional programs detectable in cord blood at birth.

### Discordant twins exhibit coordinated transcriptional reprogramming with suppressed immune signaling and enhanced erythroid transcriptional programs

To define transcriptional differences associated with sFGR, we performed differential gene expression analysis comparing twins from sFGR pairs with concordant controls (Figure. 2A and 2B), identifying 702 upregulated and 276 downregulated genes relative to concordant twins. Genes downregulated in both Disc-N and Disc-sFGR twins were strongly enriched for immune and inflammatory pathways, including regulation of innate immune responses, cytokine production, TNF and IL-6 signaling, and responses to biotic stimuli (Figure. 2C). These results indicate that the altered intrauterine environment characteristic of sFGR pregnancies is associated with systemic suppression of immune-related transcriptional programs in neonatal cord blood at birth. Genome browser views of representative immune-related loci, including *IFI44L*, *IL15*, *SIGLEC1,* and *RHBDL2*, consistently showed reduced transcriptional signals in discordant twins compared with concordant controls, supporting immune suppression at the locus level (Figure. 2D).

**Figure 2.**
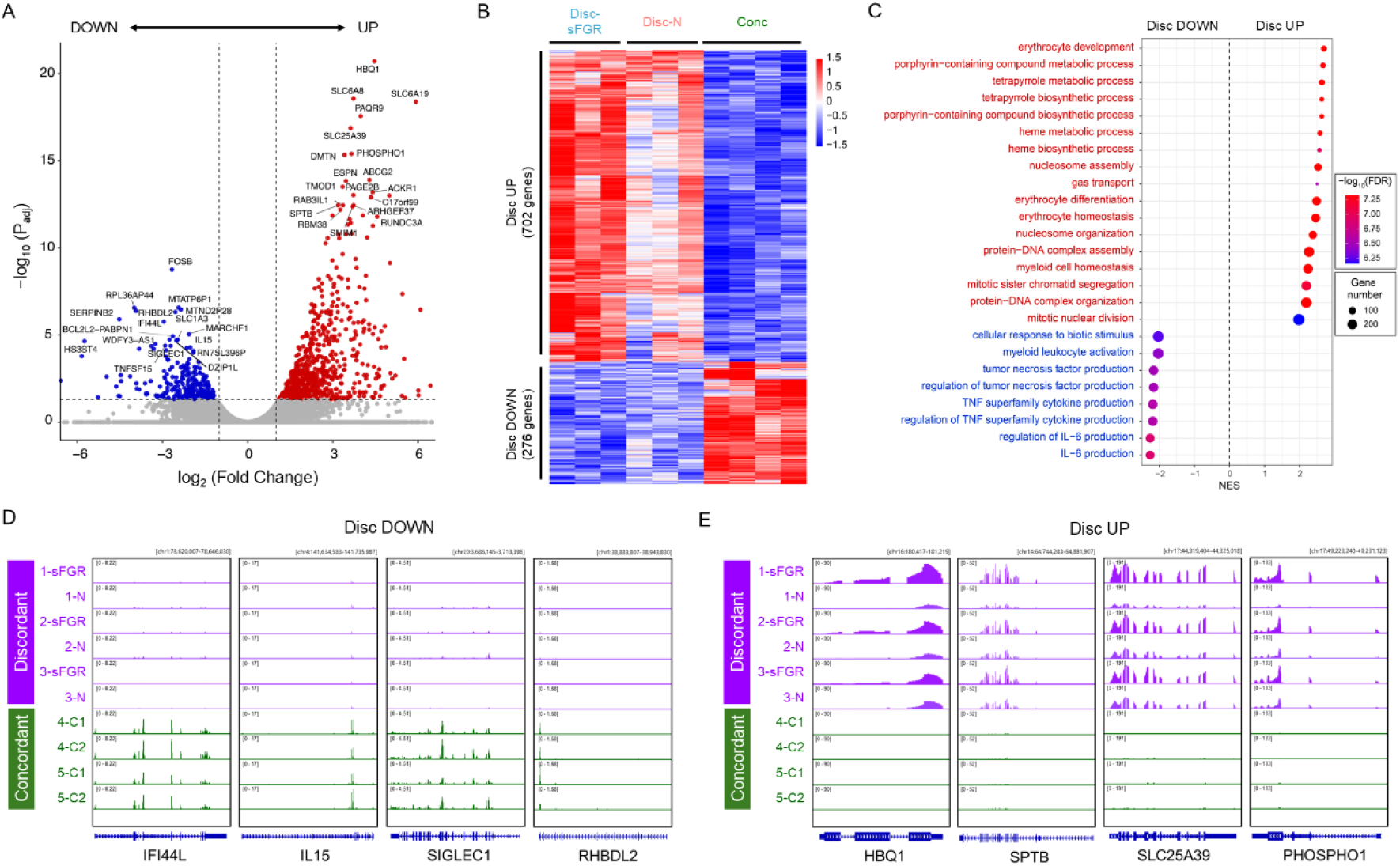
Transcriptomic signatures of discordant twins reveal enhanced erythroid programs and attenuated immune responses. (A-B) Volcano plot (A) and heatmap (B) showing differentially expressed genes between twins from discordant pairs and concordant controls. Genes meeting significance criteria (adjusted p- value < 0.05 and |log_2_(Fold Change)| > 1) are highlighted. (C) Gene set enrichment analysis (GSEA; gseGO, Biological Process) performed on the full ranked transcriptome comparing discordant twin pairs and concordant controls using pair- collapsed counts. Dot size represents gene set size; color indicates −log_10_(FDR). Positive normalized enrichment score (NES) denotes enrichment in discordant twins; negative indicates enrichment in concordant twins. (D) Genome browser tracks of representative immune-related genes (*IFI44L*, *IL15*, *SIGLEC1*, and *RHBDL2*) showing reduced transcriptional signal across discordant twins (purple) compared with concordant controls (green). (E) Genome browser tracks of representative erythroid-associated genes (*HBQ1*, *SPTB*, *SLC25A39*, and *PHOSPHO1*) showing increased transcriptional signal across discordant twins (purple) relative to concordant controls (green).

In contrast, genes upregulated in discordant twins were strongly enriched for erythroid- and chromosome-associated biological processes, including erythrocyte development and differentiation, heme and porphyrin biosynthesis, nucleosome assembly, chromatin organization, and mitotic chromosome segregation (Figure. 2C). This transcriptional signature is consistent with enhanced erythroid-associated gene activity, a response that has been linked in prior studies to fetal growth restriction and relative intrauterine hypoxic stress [37, 38]. The concomitant enrichment of chromosome- and nucleosome-related programs likely reflects increased proliferative and chromatin remodeling demands associated with erythroid lineage expansion. Correspondingly, genome browser tracks of representative loci, including *HBQ1*, *SPTB*, *SLC25A39*, and *PHOSPHO1*, showed increased transcriptional signals in discordant twins relative to concordant controls (Figure. 2E).

Together, these findings indicate coordinated hematopoietic reprogramming across discordant twin pairs, characterized by enhanced expression of erythroid-associated genes alongside suppression of innate immune and inflammatory signaling pathways. This transcriptional pattern is consistent with an adaptive response to relative intrauterine hypoxia, suggesting that developmental resources may be preferentially allocated toward oxygen transport rather than immune activation at birth.

### sFGR twins exhibit chromatin-biased transcriptional programs with selective immune rewiring

Although both co-twins within discordant pairs displayed transcriptomic divergence from concordant controls (Figure. 2), intra-pair birthweight discordance is inherently directional. We therefore performed a direct intra-pair comparison between Disc-sFGR and Disc-N twins to define transcriptional programs specifically associated with relative growth restriction. This paired analysis revealed a markedly asymmetric transcriptional response, with 627 genes upregulated and 78 genes downregulated in Disc-sFGR twins (Figure. 3A and 3B). Gene set enrichment analysis demonstrated the predominant signal in Disc-sFGR twins to be enrichment of chromosome- and cell cycle–associated biological processes, including nucleosome assembly, chromatin organization, chromosome condensation, sister chromatid segregation, mitotic nuclear division, and spindle assembly (Figure. 3C). These findings indicate that chromatin remodeling and proliferative programs represent the dominant transcriptional axis distinguishing growth-restricted twins from their genetically identical co-twins.

**Figure 3.**
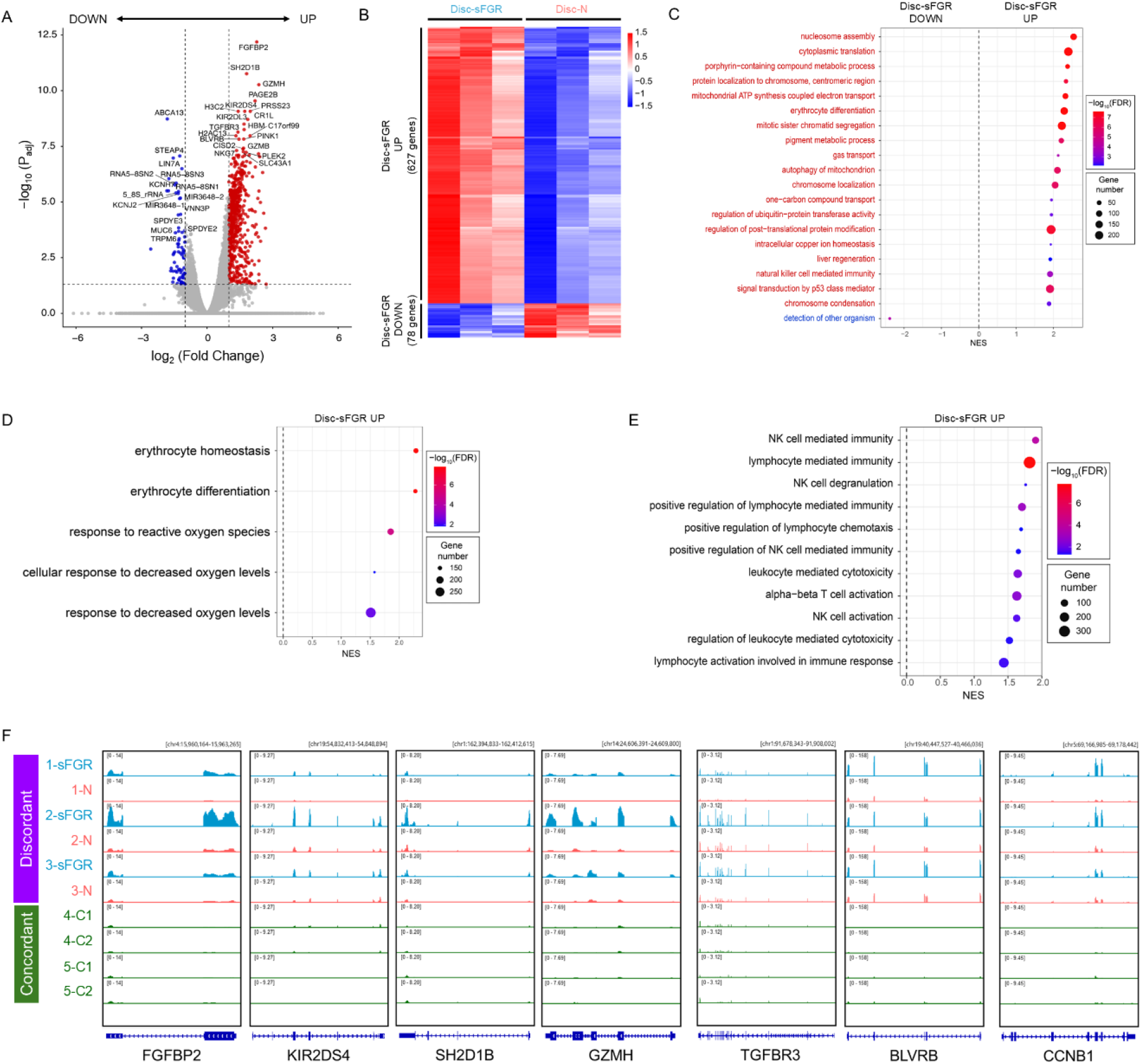
Intra-pair transcriptional asymmetry within discordant twin pairs reveals polarized states in sFGR twins. (A-B) Volcano plot (A) and heatmap (B) showing differentially expressed genes between Disc- sFGR and Disc-N twins using a paired design. Genes meeting significance criteria (adjusted p-value < 0.05 and |log_2_(Fold Change)| > 1) are highlighted. Red indicates upregulation in Disc- sFGR; blue indicates downregulation in Disc-sFGR. (C) Gene set enrichment analysis (GSEA; gseGO, Biological Process) based on the full ranked transcriptome comparing Disc-sFGR and Disc-N twins. Shown are the top enriched pathways. Dot size represents gene set size; color indicates −log_10_(FDR). Positive NES denotes enrichment in Disc-sFGR. (D) GSEA results restricted to hypoxia and oxygen response–related gene ontology terms, demonstrating stronger enrichment of hypoxia-responsive pathways in Disc-sFGR twins. (E) GSEA results restricted to immune-related gene ontology terms, showing selective enrichment of NK cell–mediated immunity and lymphocyte-associated pathways in Disc-sFGR twins. (F) Genome browser tracks of representative loci illustrating NK-associated (*FGFBP2*, *KIR2DS4*, *SH2D1B*, and *TGFBR3*), erythroid-associated (*BLVRB*) and cell cycle–associated (*CCNB1*) transcriptional activity in Disc-sFGR (blue) relative to Disc-N (red) and concordant controls (green).

In parallel, Disc-sFGR twins exhibited upregulation of bioenergetic and synthetic processes, including mitochondrial ATP synthesis coupled to electron transport and cytoplasmic translation (Figure. 3C). Gene Ontology terms related to erythroid differentiation were also strongly enriched in Disc-sFGR twins relative to Disc-N co-twins (Figure. 3C). Given that erythroid expansion is a hallmark of hypoxic adaptation, we further examined hypoxia-related pathways and observed significant enrichment of genes associated with response to decreased oxygen levels in Disc-sFGR twins (Figure. 3D), consistent with a chronic hypoxia-responsive transcriptional state.

While chromatin and proliferative programs constituted the primary axis of separation, pathway-level analysis also identified a selective shift toward natural killer (NK) cell-mediated immunity in Disc-sFGR twins (Figure. 3C). Specifically, pathways associated with NK cell activation and leukocyte-mediated cytotoxicity were significantly enriched (Figure. 3E). Core cytotoxic effector genes and NK-associated receptors, including *FGFBP2*, *KIR2DS4, SH2D1B*, and *GZMH*, were among the most strongly induced transcripts (Figure. 3A-3F).

Taken together, these data indicate that within discordant twin pairs, the growth-restricted twin exhibits a distinct and highly polarized transcriptional state. This state is characterized by coordinated activation of chromatin organization and proliferative activation alongside enhancement of erythroid-associated pathways and metabolic adaptation. The selective enrichment of NK cell signatures suggests a specific immunological response linked to relative growth restriction. Collectively, these findings highlight a coordinated molecular response to the intrauterine environment in Disc-sFGR twins, providing insights into the transcriptional features associated with fetal growth restriction.

### Myeloid programs are attenuated with concurrent activation of lymphoid and erythroid signatures in sFGR twins

Although twins from discordant pairs exhibited overall suppression of immune response genes, Disc-sFGR twins showed relatively higher expression of NK cell cytotoxic transcripts compared with Disc-N co-twins (Figure. 3E). This pattern suggests that co-twins within discordant pairs undergo distinct forms of immune remodeling. To further delineate these differences, we quantified immune cell type-specific transcriptional programs by calculating module scores based on curated marker gene sets (Figure. 4A and 4B). This analysis uncovered a structured and non-uniform reorganization of immune-associated transcriptional programs across Disc-sFGR, Disc-N, and concordant control twins. Because module scores summarize coordinated expression of lineage-specific marker genes, they likely reflect combined effects of altered cellular composition and changes in cellular transcriptional state. Module score analysis showed that Disc-sFGR twins exhibited increased erythroid/heme, cell cycle, and oxidative phosphorylation programs, accompanied by increased activity of NK cell-, T cell-, and hematopoietic stem and progenitor cell (HSPC)-associated transcriptional modules compared with both Disc-N co-twins and concordant controls (Figure. 4A-4C). In contrast, transcriptional programs associated with monocytes, neutrophils, dendritic cells, B cells, and interferon responses were consistently downregulated in both Disc-sFGR and Disc-N twins relative to concordant controls (Figure. 4C), indicating a broad attenuation of myeloid- and innate immune-associated programs across discordant pregnancies.

**Figure 4.**
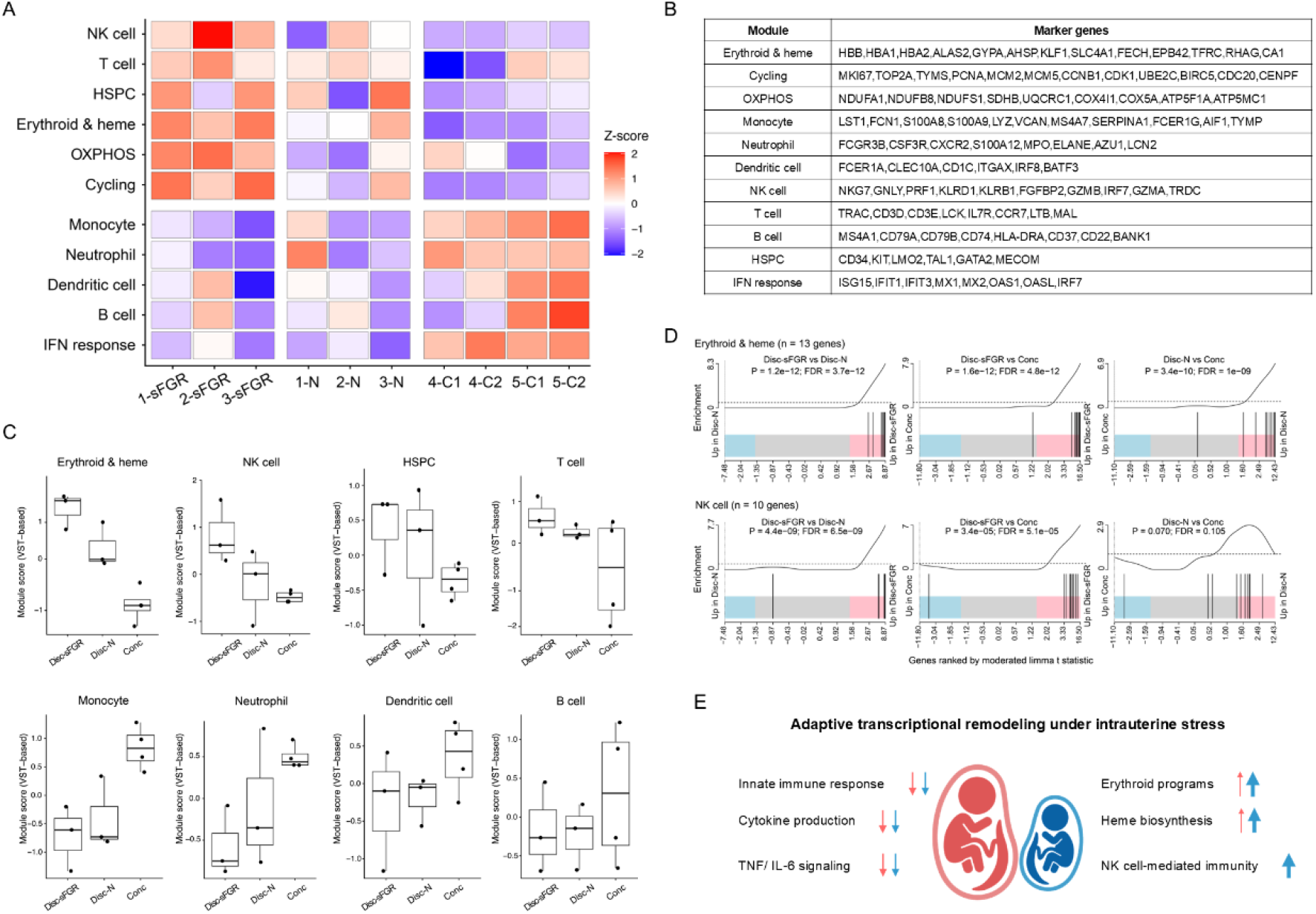
Module score analysis highlights structured remodeling of immune transcriptional programs across discordant and concordant twin pairs. (A-B) Heatmap (A) of immune- and hematopoietic-lineage module scores across individual samples from Disc-sFGR, and Disc-N, and concordant controls twins, calculated using the curated marker gene sets listed in (B). (C) Module scores for major immune and hematopoietic lineages, including NK cells, T cells, hematopoietic stem and progenitor cells (HSPCs), erythroid/heme, monocytes, neutrophils, dendritic cells, B cells, and interferon response pathways. (D) Barcode plots depicting the rank positions of erythroid/heme (top) and NK cell (bottom) marker genes within transcriptomes ranked by moderated limma t-statistic for each pairwise comparison. Vertical bars mark individual marker gene positions, and the curve above each plot shows relative enrichment along the ranked list. (E) Schematic summary of the transcriptional remodeling associated with prenatal growth discordance, highlighting shared attenuation of innate immune and inflammatory signaling (cytokine production, TNF/IL-6) alongside selective enrichment of erythroid, heme, and NK cell-mediated programs in Disc-sFGR twins.

To visualize the coordinated behavior of genes within each predefined lineage-specific module, marker genes were mapped onto the ranked transcriptome according to differential expression between groups (Figure. 4D). Erythroid/heme marker genes were strongly concentrated among the most highly upregulated genes in Disc-sFGR in all three comparisons, demonstrating concerted activation of the erythroid transcriptional program. Likewise, NK cell marker genes clustered toward the upregulated end of the ranked transcriptome in Disc-sFGR compared with both Disc-N co-twins and concordant controls, indicating coordinated activation of the cytotoxic lymphoid program.

Together, these findings indicate that fetal growth restriction is associated not with uniform immune suppression but with selective immune remodeling. While myeloid- and interferon-associated programs are broadly dampened across discordant twins, sFGR twins display coordinated activation of erythroid and cytotoxic lymphoid transcriptional programs (Figure. 4E). This imbalance suggests a skewing toward erythroid- and lymphoid-associated immune programs in Disc-sFGR neonates within the broader context of attenuated innate immune function.

### Genome-wide DNA methylation profiling identifies immune-associated differentially methylated regions in sFGR twins

Given the coordinated transcriptional activation of NK cell–associated and hypoxia-responsive transcription programs in Disc-sFGR twins, we next examined whether these alterations were accompanied by corresponding epigenetic changes at immune-related loci. We evaluated the sequencing quality of all samples before performing differential methylation analysis. Mapping efficiency ranged from 81.2% to 82.4%, and bisulfite conversion rates were consistently ≥99.3% across all samples (Figure. S2A). We identified distinct differentially methylated regions (DMRs) between the discordant twins, including 1047 hypomethylated and 1093 hypermethylated loci in Disc-sFGR twins (Figure. 5A). Principal component analysis restricted to CpGs within the identified DMRs (n=4,140; coverage ≥5 in all samples) demonstrated clear separation of Disc-sFGR and Disc-N twins along PC1 (50.8% of variance), supporting the robustness of the identified DMRs (Figure. S2B). Interestingly, hypomethylated regions in Disc- sFGR twins were enriched for pathways related to lymphocyte differentiation, leukocyte activation and T cell differentiation (Figure. 5B and S2C) consistent with the transcriptional signatures observed in the RNA-seq analysis.

**Figure 5.**
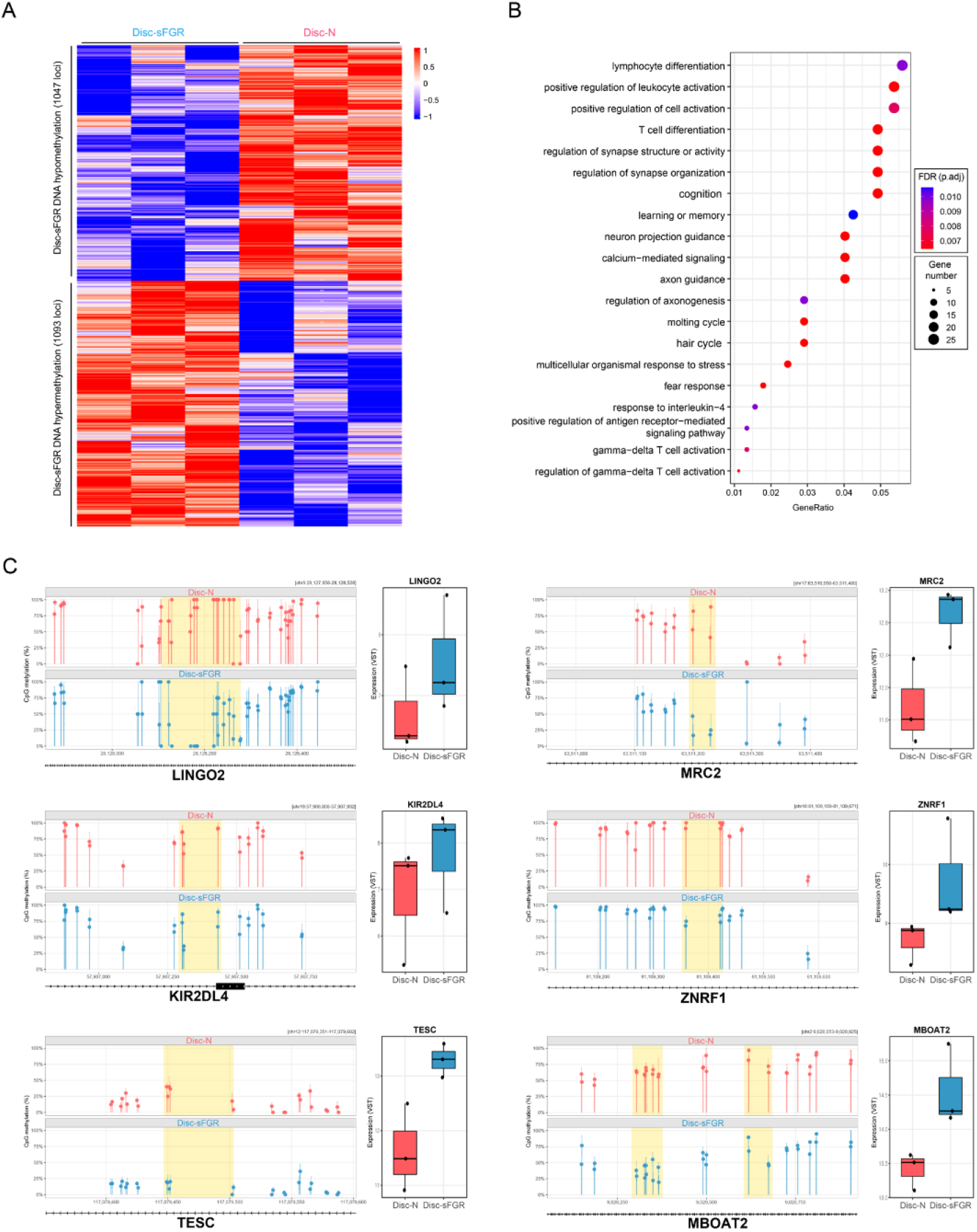
Differential DNA methylation and concordant transcriptional changes in Disc- sFGR twins. (A) Heatmap showing differentially methylated regions (DMRs) between Disc-sFGR and Disc-N twins. DMRs were called by aggregating contiguous CpG loci with Wald test p < 0.1, requiring a minimum length of 30 bp, at least 2 CpGs, and |Δmethylation| ≥ 20%. A total of 1,047 hypomethylated and 1,093 hypermethylated loci were identified in Disc-sFGR twins. (B) Gene Ontology (GO) enrichment analysis of genes associated with hypomethylated DMRs in Disc-sFGR twins. Enriched biological processes include lymphocyte differentiation, leukocyte activation, T cell differentiation, and immune-related signaling pathways. Dot size represents gene count and color indicates adjusted p-value. (C) Lollipop plots and RNA-seq expression profiles of genes exhibiting concordant hypomethylation and transcriptional upregulation in Disc-sFGR twins, identified by overlap between DMR-associated genes and DEGs (adjusted p-value < 0.05 and |log2(Fold Change)| > 1) within discordant twin analysis. For each gene, the left panel displays CpG methylation levels (%) in Disc-N (red) and Disc-sFGR (blue) twins, with yellow shaded regions indicating DMRs. The right panel shows VST-normalized RNA-seq expression values for each group.

Integrative analysis further identified concordant hypomethylation and upregulation of gene expression at several loci in Disc-sFGR twins (Figure. S2D), including *LINGO2, MRC2, KIR2DL4, ZNRF1, TESC,* and *MBOAT2*, suggesting a potential link between epigenetic remodeling and transcriptional changes. These genes have known or emerging roles in immune regulation and stress adaptation. *KIR2DL4* is an NK cell receptor involved in maternal–fetal immune signaling, while *TESC* has been implicated in hematopoietic and T-cell differentiation. *ZNRF1, MRC2, MBOAT2,* and *LINGO2* are involved in inflammatory signaling, immune reconstitution, and metabolic adaptation, and these factors collectively support selective immune rewiring in growth-restricted twins.

Moreover, several loci exhibited differential promoter-proximal methylation without a clear association with corresponding transcriptional changes. For example, *ATP7A*, a hypoxia-responsive copper transporter, and *LIG4*, a key regulator of lymphocyte development via V(D)J recombination, showed distinct methylation patterns between the discordant twins (Figure. S2E). This suggests the existence of epigenetic variations that may reflect latent regulatory potential, although they do not lead to direct and immediate transcriptional divergence. Although direct mechanical linkages have not been established at all gene loci, these data collectively show a coordinated pattern of epigenetic remodeling at immune and hypoxia-related gene loci in sFGR twins. This epigenetic landscape may represent an early imprint of intrauterine environmental conditions, potentially contributing to long-term immune programming and disease susceptibility. To assess whether these transcriptional and DNA methylation differences could be attributed to changes in cell-type composition rather than cell-intrinsic regulatory alterations, we performed EpiDISH-based deconvolution of whole-genome bisulfite sequencing (WGBS) data using a cord-blood-specific reference panel (Figure. S3A). No significant differences were observed in the putative cell type composition of the Disc-N and Disc-sFGR twins (all p ≥ 0.06, paired t-test, n = 3 pairs) (Figure. S3B). Although these estimates were not independently validated by scRNA-seq or flow cytometry, the findings argue against cell composition as the primary driver of the observed molecular differences and instead support cell-intrinsic epigenetic reprogramming in response to the adverse intrauterine environment experienced by growth-restricted twins.

## Discussion

In this study, we leveraged a rare cohort of MZ-DCDA twins to define neonatal transcriptional programs associated with prenatal growth discordance while rigorously controlling for genetic background. By profiling umbilical cord blood at birth, we captured immune and hematopoietic transcriptional states shaped by intrauterine conditions before postnatal exposures could confound early-life programming. This design enabled us to link transcriptional and epigenetic differences to divergent intrauterine environments within genetically identical individuals. When comparing twins from sFGR pairs with concordant-growth controls, we observed structured immune remodeling rather than uniform immune suppression. Namely, transcriptional programs linked to myeloid and antigen-presenting compartments—including monocytes, neutrophils, and dendritic cells—as well as interferon signaling were attenuated across discordant twins, whereas erythroid differentiation, heme biosynthesis, and chromatin-associated pathways were enriched. These findings suggest that fetal growth restriction is accompanied by a redistribution of hematopoietic priorities at birth, characterized by relative dampening of innate myeloid and interferon programs alongside enhanced erythroid activity.

Direct intra-pair comparison between discordant-sFGR (Disc-sFGR) and discordant-normal (Disc-N) co-twins further revealed a pronounced asymmetry. In Disc-sFGR twins, the dominant transcriptional axis reflected enrichment of erythroid and gas transport signatures, indicating a proliferative and bioenergetically activated hematopoietic state. Because erythroid expansion is closely linked to oxygen availability, the enrichment of hypoxia-responsive gene ontology terms supports a model of adaptive hematopoietic remodeling under intrauterine stress. Although hypoxia was not directly measured, the coordinated activation of erythroid, mitochondrial, and chromatin-related programs is consistent with a compensatory adaptation rather than a primary cause of growth restriction.

Notably, immune alterations within discordant pairs were selective. Among the pathways enriched in Disc-sFGR twins, a cytotoxic lymphoid signature, attributable to natural killer (NK) and/or cytotoxic T cells, emerged as a reproducible signal, with several core cytotoxic effectors and NK receptors among the most strongly induced transcripts. Because these cytotoxic effector genes are largely shared between NK and CD8 T cells, the precise lineage of this expanded compartment cannot be resolved from bulk data. This induction occurred in the broader context of reduced myeloid and interferon signatures in discordant twins relative to concordant controls. Together, these findings suggest a developmental “lineage tilt” in fetal hematopoiesis— specifically an expansion of erythroid and cytotoxic lymphoid programs. Such skewing may contribute to the vulnerability of sFGR neonates to infection, as suppression of antigen-presenting and innate myeloid pathways could compromise early immune responsiveness. In particular, since circulating monocytes serve as essential precursors for tissue-resident macrophages, the observed transcriptomic suppression suggests a reduction in myeloid programming at birth. Functional defects in conserved myeloid processes, such as those governing cellular homeostasis and autophagy, have been shown to impair long-term immune resilience and heighten inflammatory risks [39]. Conversely, enhanced NK-associated programs may reflect stress-induced maturation of the innate lymphoid compartment. Consistent with this, recent studies have demonstrated that NK cell activity is precisely fine-tuned by extrinsic signals to coordinate early-life immune responsiveness [40], highlighting the functional plasticity of NK cells within specific stress environments [41]. Integrative DNA methylation analysis further revealed that the transcriptional reprogramming observed in Disc-sFGR twins is accompanied by coordinated epigenetic remodeling. Hypomethylated DMRs in Disc-sFGR twins were enriched for immune-related pathways, including lymphocyte differentiation and leukocyte activation, consistent with the transcriptional signatures identified in RNA-seq analysis. At select loci, concordant hypomethylation and transcriptional upregulation were observed, suggesting that epigenetic changes may contribute to the selective immune remodeling in growth-restricted neonates. These findings raise the possibility that intrauterine growth restriction leaves durable epigenetic imprints on immune-associated loci, potentially influencing immune programming beyond the neonatal period.

Our study design is central to this interpretation. By focusing on MZ-DCDA twins, which develop within independent placentas and amniotic sacs, we minimized shared vascular confounders while preserving identical genetic backgrounds. Unlike murine models of uterine artery ligation or nutritional restriction, which are valuable but occur over a compressed gestational timeline, this human model captures chronic, naturally occurring placental divergence across months of development. To our knowledge, genome-wide transcriptomic and epigenomic profiling of cord blood from MZ-DCDA twins discordant for birthweight has not been previously reported. As only one to three such eligible DCDA monozygotic twin cases are identified annually at our institution, this cohort represents a uniquely informative model for studying intrauterine environmental programming.

These findings extend prior epidemiologic observations linking birthweight to immune vulnerability and long-term disease risk. Although much of the early-life programming literature has focused on metabolic and cardiovascular outcomes, our data provide molecular evidence that immune lineage allocation and hematopoietic priorities are already structured at birth in association with prenatal growth conditions. Consistent with this view, birthweight is a well-established predictor of neonatal immune function and long-term health outcomes. Low-birthweight infants exhibit increased susceptibility to infection and mortality, which has been associated with downregulation of interferon-signature genes and impaired hematopoietic stem cell differentiation [42, 43]. Conversely, high birthweight has been linked to an elevated risk of acute lymphoblastic leukemia [44]. Together, these observations suggest that the neonatal immune system may adopt a compensatory but potentially imbalanced “stress-primed” configuration in the context of intrauterine growth restriction.

Several limitations should be noted. Because profiling was performed on bulk umbilical cord blood buffy coat, the observed transcriptional signatures likely reflect a combination of changes in cellular composition and cell-intrinsic regulatory programs, making it difficult to distinguish between these effects. In particular, the relative attenuation of myeloid and interferon pathways and enrichment of cytotoxic lymphoid signatures may partly arise from hematologic alterations associated with fetal growth restriction. Furthermore, key cytotoxic transcripts are shared across multiple lymphoid populations, and highly polymorphic *KIR* genes are challenging to quantify accurately using short-read sequencing, limiting precise identification of the affected cell populations. Although concordant DNA methylation and expression changes were observed at selected loci, a direct mechanistic relationship was not established genome wide. Future studies integrating single-cell and epigenomic profiling will be required to define the precise identity of the expanded cytotoxic lymphoid compartment and determine whether fetal growth restriction primarily alters cellular composition, functional state, or both. These approaches will also help clarify the regulatory basis and long-term consequences of the transcriptional and epigenetic alterations observed at birth. Nevertheless, the consistent pathway-level changes across twin pairs support the conclusion that the intrauterine environment can shape neonatal immune and hematopoietic programming independently of genetic variation.

## Conclusions

In conclusion, prenatal growth discordance in MZ-DCDA twins is associated with coordinated remodeling of neonatal hematopoiesis, characterized by activation of chromatin and proliferative pathways, augmented erythroid and hypoxia-associated programs, selective enrichment of cytotoxic lymphoid (NK/T cell), and attenuation of myeloid and interferon-related pathways. Integrative DNA methylation analysis further identified concordant epigenetic remodeling at immune-associated loci, suggesting that these transcriptional programs are accompanied by durable epigenetic imprints shaped by intrauterine conditions. These findings provide evidence from a genetically controlled human twin model that intrauterine growth conditions are associated with both transcriptional and epigenetic dimensions of immune lineage architecture at the time of birth.

## List of abbreviations

BH: Benjamini-Hochberg
Conc: Concordant control twin pairs
DCDA: Dichorionic–diamniotic
DEG: Differentially expressed gene
Disc-N: Normal-birthweight co-twin from a selective fetal growth restriction pair
Disc-sFGR: Growth-restricted co-twin from a selective fetal growth restriction pair
DML: Differentially methylated locus
DMR: Differentially methylated region
GO: Gene ontology
GSEA: Gene set enrichment analysis
HSPC: Hematopoietic stem and progenitor cell
MZ: Monozygotic
MZ-DCDA: Monozygotic dichorionic–diamniotic
NK: Natural killer
PCA: Principal component analysis
RNA-seq: RNA sequencing
RPC: robust partial correlation
sFGR: Selective fetal growth restriction
STR: short tandem repeat
VST: Variance-stabilizing transformation
WGBS: Whole-genome bisulfite sequencing

## Declarations

### Ethics statement (Ethics approval and consent to participate)

The Institutional Review Board of Seoul National University Hospital approved this study [1909-054-1063]. Written consent for the use of clinical information and biospecimens was obtained from the participants’ parents. Additional consent for further research was waived by the IRB. The study was conducted in accordance with the ethical principles of the Declaration of Helsinki.

## Consent for publication

Not applicable

## Data availability

The datasets used in this research are accessible by the corresponding authors upon reasonable request.

## Competing interests

The authors declare no competing interests.

## Funding

This work was supported by the Korean government (MSIT) (NRF2021R1C1C1013220, RS- 2022-NR070833, RS-2023-00223069, RS-2023-00211612, RS-2024-00411768, RS- 2025024523684, RS-2025-02217909), the BK21 Four Biomedical Science Program, and the SNUH Kun-hee Lee Child Cancer and Rare Disease Project Foundation (22B-001-0100). Additional support was provided by the Research Resettlement Fund for New Faculty at SNU, the Creative-Pioneering Researchers Program (SNU), the SNUBH–SNU Medicine Collaborative Research Fund (16-2026-0001), Seoul National University College of Medicine (550-20250081), the AI-Bio Research Grant (SNU), the Doosan Yonkang Foundation (SNUH-30-2024-0440), and NAVER (SNUH-3720242170). This study was also funded by SNUH Research Fund (1120215040) and by the National Research Foundation of Korea (NRF) grant funded by the Korea government (MSIT) (RS-2025-00558188). The Biospecimens and data used in this study were provided by the Biobank of Seoul National University Hospital, a member of Korea Biobank Network (KBN4_A03). This research was also supported by the Suh Kyungbae Foundation (no. SUHF-24010031 to J.-Y.L.); by the National Research Foundation of Korea funded by the Korean Government (no. RS-2023-00217798 and no. RS-2024-00349044 to J.Y.L).

## Authors’ contributions

H.K., S.K., S.M.L. and C.-H.L. conceptualized and designed the study. H.K. and S.K. conducted the experiments and mainly analyzed transcriptome and DNA methylation datasets. S.K., J.H.K., C.-W.P., J.S.P., J.K.J. and S.M.L. provided clinical data and prepared buffy coat samples from concordant and discordant twin pairs. J.-Y.L., D.L. and C.-H.L. further analyzed and validated the transcriptomic and epigenomic datasets. All authors read and approved the final manuscript.

## Acknowledgements

We thank members of the Lee laboratory for valuable discussions.

## Additional files

**Additional file 1 (.pdf): Supplementary figures.** Figures S1–S3 showing RNA-seq quality-control metrics, supplementary DNA methylation analyses, and EpiDISH-based cell-type deconvolution.

## Notes

### Competing Interest Statement

The authors have declared no competing interest.

## References

1. Boomsma DI: Twin, association and current "omics" studies. J Matern Fetal Neonatal Med 2013, 26 Suppl 2:9–12.

2. Turrina S, Bortoletto E, Giannini G, De Leo D: Monozygotic twins: Identical or distinguishable for science and law? Med Sci Law 2021, 61:62–66.

3. Wong AH, Gottesman, II, Petronis A: Phenotypic differences in genetically identical organisms: the epigenetic perspective. Hum Mol Genet 2005, 14 Spec No 1:R11–18.

4. Weidman JR, Dolinoy DC, Murphy SK, Jirtle RL: Cancer susceptibility: epigenetic manifestation of environmental exposures. Cancer J 2007, 13:9– 16.

5. Nyberg DA, Filly RA, Golbus MS, Stephens JD: Entangled umbilical cords: a sign of monoamniotic twins. J Ultrasound Med 1984, 3:29–32.

6. Derom C, Thiery E, Rutten BPF, Peeters H, Gielen M, Bijnens E, Vlietinck R, Weyers S: The East Flanders Prospective Twin Survey (EFPTS): 55 Years Later. Twin Res Hum Genet 2019, 22:454–459.

7. Corey LA, Nance WE, Kang KW, Christian JC: Effects of type of placentation on birthweight and its variability in monozygotic and dizygotic twins. Acta Genet Med Gemellol (Roma) 1979, 28:41–50.

8. Blickstein I: Monochorionicity in perspective. Ultrasound Obstet Gynecol 2006, 27:235–238.

9. Salafia CM, Maas E: The twin placenta: framework for gross analysis in fetal origins of adult disease initiatives. Paediatr Perinat Epidemiol 2005, 19 Suppl 1:23–31.

10. Machin G, Still K, Lalani T: Correlations of placental vascular anatomy and clinical outcomes in 69 monochorionic twin pregnancies. Am J Med Genet 1996, 61:229–236.

11. Race JP, Townsend GC, Hughes TE: Chorion type, birthweight discordance and tooth-size variability in Australian monozygotic twins. Twin Res Hum Genet 2006, 9:285–291.

12. Loos RJ, Derom C, Derom R, Vlietinck R: Determinants of birthweight and intrauterine growth in liveborn twins. Paediatr Perinat Epidemiol 2005, 19 Suppl 1:15–22.

13. Weiner E, Barber E, Feldstein O, Dekalo A, Schreiber L, Bar J, Kovo M: Placental Histopathology Differences and Neonatal Outcome in Dichorionic-Diamniotic as Compared to Monochorionic-Diamniotic Twin Pregnancies. Reprod Sci 2018, 25:1067–1072.

14. Bakwin H: Body-weight Regulation in Twins. Developmental Medicine & Child Neurology 1973, 15:178–183.

15. Clemetson C: THE DIFFERENCE IN BIRTH WEIGHT OF HUMAN TWINS: Twin Blood Studies: I—Oxygen Analysis of Umbilical Cord Blood. BJOG: An International Journal of Obstetrics & Gynaecology 1956, 63:1–8.

16. Lee K, Hur J, Yoo J: Twin weight discordance and maternal weight gain in twin pregnancies. International Journal of Gynecology & Obstetrics 2007, 96:176–180.

17. Conley D, Strully KW, Bennett NG: Twin differences in birth weight: the effects of genotype and prenatal environment on neonatal and post- neonatal mortality. Economics & Human Biology 2006, 4:151–183.

18. Canpolat FE, Çekmez F, Sarici SÜ, Korkmaz A, Yurdakok M: Insulin-like growth factor-1 levels in twins and its correlation with discordance. Twin Research and Human Genetics 2011, 14:94–97.

19. Lim SY, Shin SH, Yang HJ, Park SG, Kim E-K, Kim H-S, Jun JK: Neonatal and developmental outcomes of very preterm twins according to the chorionicity and weight discordance. Scientific Reports 2023, 13:6784.

20. Magnus P, Berg K, Bjerkedal T: No significant difference in birth weight for offspring of birth weight discordant monozygotic female twins. Early human development 1985, 12:55–59.

21. Heijmans BT, Tobi EW, Stein AD, Putter H, Blauw GJ, Susser ES, Slagboom PE, Lumey LH: Persistent epigenetic differences associated with prenatal exposure to famine in humans. Proc Natl Acad Sci U S A 2008, 105:17046– 17049.

22. Tobi EW, Goeman JJ, Monajemi R, Gu H, Putter H, Zhang Y, Slieker RC, Stok AP, Thijssen PE, Muller F, et al: DNA methylation signatures link prenatal famine exposure to growth and metabolism. Nat Commun 2014, 5:5592.

23. Martino D, Kresoje N, Amenyogbe N, Ben-Othman R, Cai B, Lo M, Idoko O, Odumade OA, Falsafi R, Blimkie TM, et al: DNA Methylation signatures underpinning blood neutrophil to lymphocyte ratio during first week of human life. Nat Commun 2024, 15:8167.

24. Curado J, Sileo F, Bhide A, Thilaganathan B, Khalil A: Early- and late-onset selective fetal growth restriction in monochorionic diamniotic twin pregnancy: natural history and diagnostic criteria. Ultrasound Obstet Gynecol 2020, 55:661–666.

25. Mazer Zumaeta A, Gil MM, Rodriguez-Fernandez M, Carretero P, Ochoa JH, Casanova MC, Molina FS: Selective Fetal Growth Restriction in Monochorionic Diamniotic Twins: Diagnosis and Management. Matern Fetal Med 2022, 4:268–275.

26. van Gemert MJ, Umur A, Tijssen JG, Ross MG: Twin-twin transfusion syndrome: etiology, severity and rational management. Curr Opin Obstet Gynecol 2001, 13:193–206.

27. Kusanovic JP, Romero R, Gotsch F, Mittal P, Erez O, Kim CJ, Hassan SS, Espinoza J, Yeo L: Discordant placental echogenicity: a novel sign of impaired placental perfusion in twin-twin transfusion syndrome? J Matern Fetal Neonatal Med 2010, 23:103–106.

28. Selmi C, Cavaciocchi F, Lleo A, Cheroni C, De Francesco R, Lombardi SA, De Santis M, Meda F, Raimondo MG, Crotti C, et al: Genome-wide analysis of DNA methylation, copy number variation, and gene expression in monozygotic twins discordant for primary biliary cirrhosis. Front Immunol 2014, 5:128.

29. Petronis A, Gottesman, II, Kan P, Kennedy JL, Basile VS, Paterson AD, Popendikyte V: Monozygotic twins exhibit numerous epigenetic differences: clues to twin discordance? Schizophr Bull 2003, 29:169–178.

30. Gordon L, Joo JH, Andronikos R, Ollikainen M, Wallace EM, Umstad MP, Permezel M, Oshlack A, Morley R, Carlin JB, et al: Expression discordance of monozygotic twins at birth: effect of intrauterine environment and a possible mechanism for fetal programming. Epigenetics 2011, 6:579–592.

31. Wadji DL, Nemoda Z, Martin-Soelch C, Booij L, Wicky C: Birth weight discordance, gene expression, and DNA methylation: A scoping review of epigenetic twin studies. PLoS One 2025, 20:e0315549.

32. Lee KA, Oh KJ, Lee SM, Kim A, Jun JK: The frequency and clinical significance of twin gestations according to zygosity and chorionicity. Twin Res Hum Genet 2010, 13:609–619.

33. Wu D, Smyth GK: Camera: a competitive gene set test accounting for inter-gene correlation. Nucleic Acids Res 2012, 40:e133.

34. Teschendorff AE, Breeze CE, Zheng SC, Beck S: A comparison of reference-based algorithms for correcting cell-type heterogeneity in Epigenome-Wide Association Studies. BMC Bioinformatics 2017, 18:105.

35. Guo X, Sulaiman M, Neumann A, Zheng SC, Cecil CAM, Teschendorff AE, Heijmans BT: Unified high-resolution immune cell fraction estimation in blood tissue from birth to old age. Genome Med 2025, 17:63.

36. Kalish RB, Chasen ST, Gupta M, Sharma G, Perni SC, Chervenak FA: First trimester prediction of growth discordance in twin gestations. Am J Obstet Gynecol 2003, 189:706–709.

37. Haase VH: Regulation of erythropoiesis by hypoxia-inducible factors. Blood Rev 2013, 27:41–53.

38. Ducsay CA, Goyal R, Pearce WJ, Wilson S, Hu XQ, Zhang L: Gestational Hypoxia and Developmental Plasticity. Physiol Rev 2018, 98:1241–1334.

39. Zhu W, Chen Z, Gao Y, Zhai C, Li X, Wang N, Fu K, Chen W, Peng J, Xu D, et al: Deficient chaperone-mediated autophagy in macrophages aggravates colitis and colitis-associated tumorigenesis in mice. Mol Cells 2025, 48:100298.

40. Ashraf MU, Kang MH, Lim YT, Yang S, Bae YS: Egr2-dependent Mo-DCs regulate immunosuppressive NK cells, promoting neutrophil-mediated protective immunity against acute Listeria infection. Mol Cells 2025, 48:100287.

41. Nathalie G, Bonamichi B, Kim J, Jeong J, Kang H, Hartland ER, Eveline E, Lee J: NK cell-activating receptor NKp46 does not participate in the development of obesity-induced inflammation and insulin resistance. Mol Cells 2024, 47:100007.

42. Singh S, Singh VK, Rai G: Identification of Differentially Expressed Hematopoiesis-associated Genes in Term Low Birth Weight Newborns by Systems Genomics Approach. Curr Genomics 2019, 20:469–482.

43. Singh VV, Chauhan SK, Rai R, Kumar A, Singh SM, Rai G: Decreased pattern recognition receptor signaling, interferon-signature, and bactericidal/permeability-increasing protein gene expression in cord blood of term low birth weight human newborns. PLoS One 2013, 8:e62845.

44. Che H, Long D, Sun Q, Wang L, Li Y: Birth Weight and Subsequent Risk of Total Leukemia and Acute Leukemia: A Systematic Review and Meta-Analysis. Front Pediatr 2021, 9:722471.

