## Supplemental Figure for "Intrauterine growth restriction is associated with adaptive hematopoietic reprogramming and selective immune rewiring in monozygotic twins"

**Supplemental Information**

**Supplemental Figures and Figure Legends**

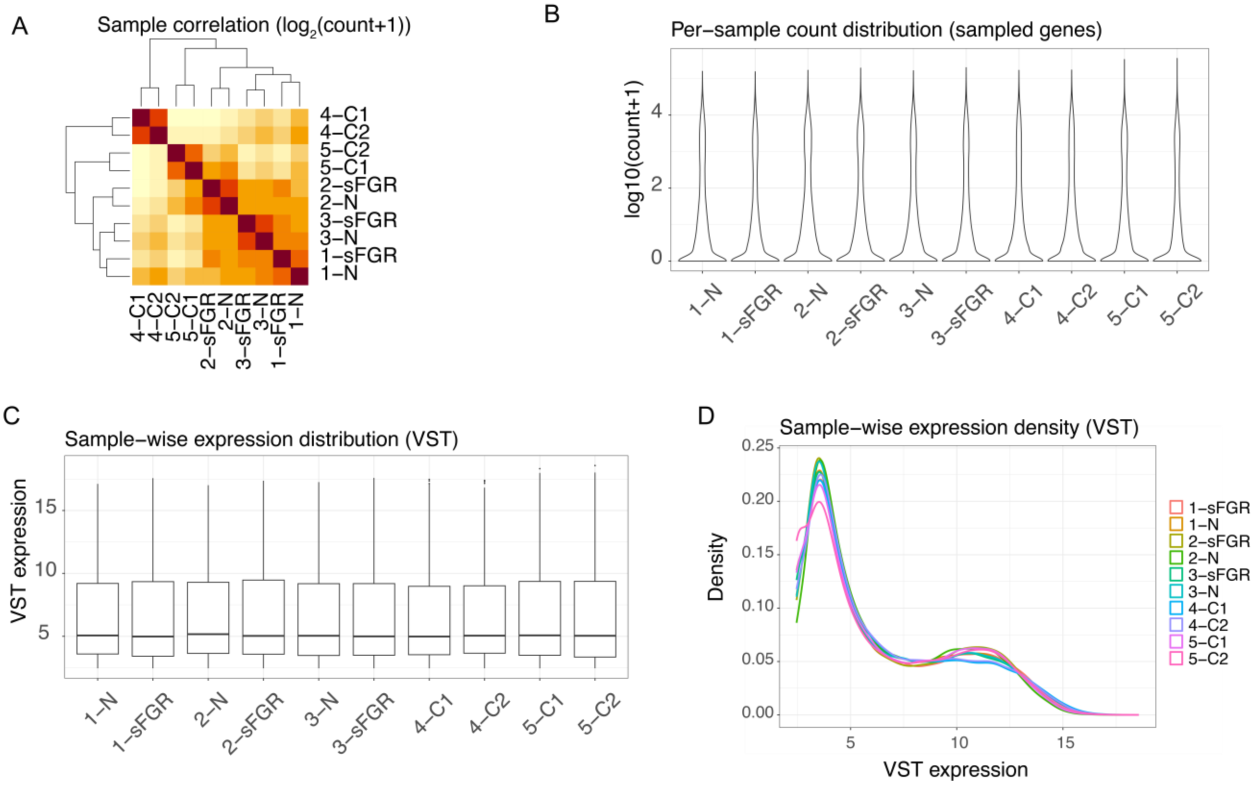

**Figure S1. Quality control assessment of RNA-seq data from frozen buffy coat–derived samples.**

(A) Sample-to-sample correlation heatmap based on $log_2$(count+1) transformed expression values, showing high overall concordance and appropriate clustering of biologically related samples.

(B) Distribution of gene expression counts ($log_{10}$[count+1]) across samples using a subsampled gene set, demonstrating comparable library complexity and dynamic range.

(C) Boxplot of variance-stabilized (VST) expression values across all samples, indicating consistent global expression distributions without evidence of technical bias.

(D) Density plot of VST-transformed expression values, showing highly overlapping profiles across samples, consistent with uniform normalization and data quality.

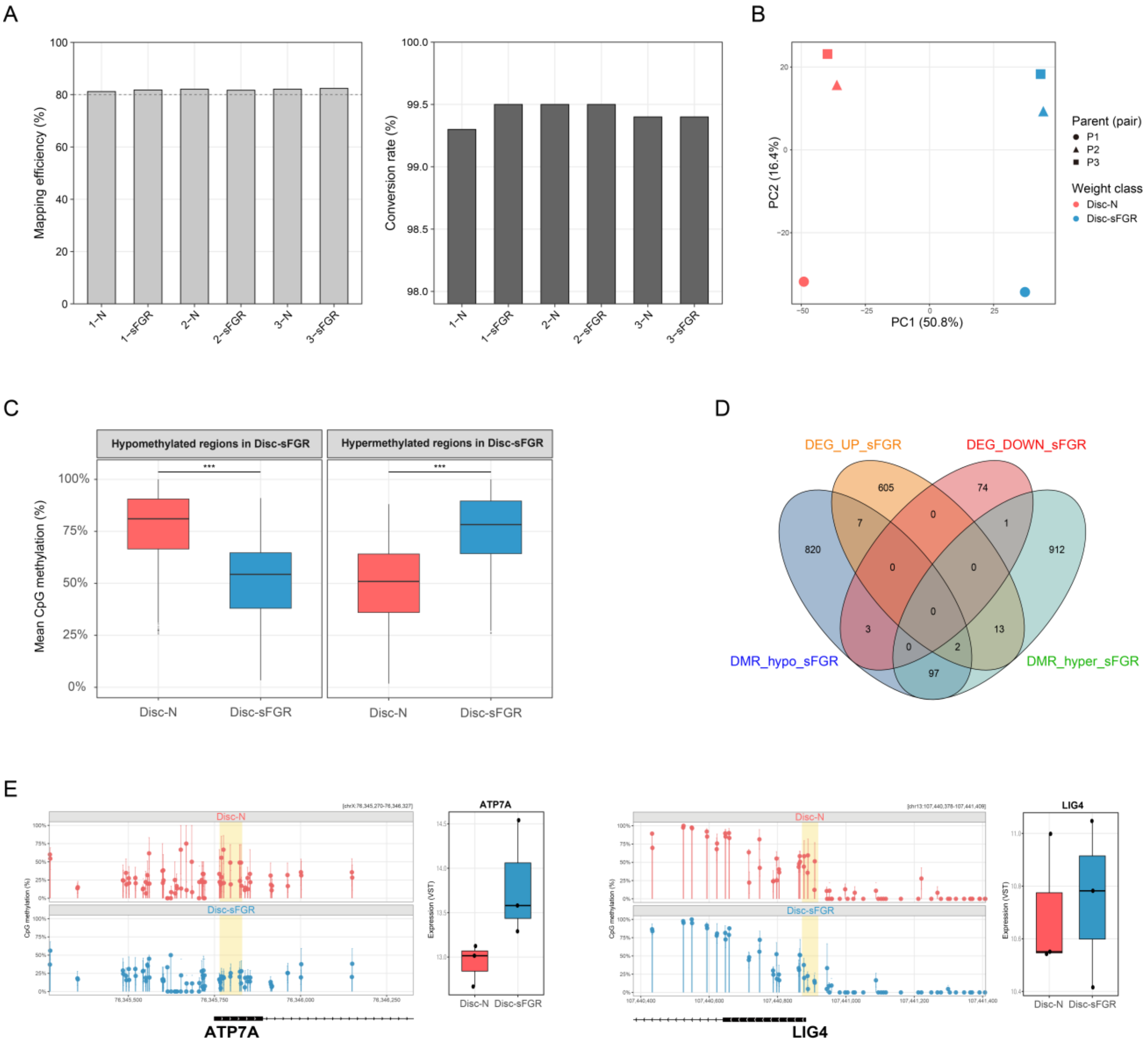

A
Mapping efficiency (%)
1-N
1-sFGR
2-N
2-sFGR
3-N
3-sFGR
Conversion rate (%)
B
PC1 (50.8%)
PC2 (16.4%)
Parent (pair)
P1
P2
P3
Weight class
Disc-N
Disc-sFGR
C
Hypomethylated regions in Disc-sFGR
Hypermethylated regions in Disc-sFGR
Mean CpG methylation (%)
***
Disc-N
Disc-sFGR
D
DEG_UP_sFGR
DEG_DOWN_sFGR
DMR_hypo_sFGR
DMR_hyper_sFGR
605
74
820
912
7
0
1
0
0
0
3
13
0
2
97
E
Disc-N
Disc-sFGR
CpG methylation (%)
ATP7A
Expression (VST)
LIG4

**Figure S2. WGBS quality control and integrative comparison in Disc-sFGR twins**

(A) Mapping efficiency (left) and bisulfite conversion rate (right) for each sample. Dashed line indicates 80% mapping efficiency. Bisulfite conversion rate was estimated from non-CpG (CHH and CHG context) methylation levels. All six samples showed mapping efficiency > 81% and bisulfite conversion rate > 99.3%.

(B) Principal component analysis (PCA) was performed on per-CpG methylation values restricted to CpGs within identified DMRs (coverage ≥ 5 in all six samples; n = 4,140 CpGs).

(C) Box plots showing mean CpG methylation levels across hypomethylated (left) and hypermethylated (right) DMRs identified in Disc-sFGR twins relative to Disc-N co-twins. Hypomethylated DMRs showed significantly lower CpG methylation in Disc-sFGR compared to Disc-N twins (median 54.3% vs. 81.0%, Wilcoxon signed-rank test, $p < 2.3 \times 10^{-173}$), while hypermethylated DMRs showed significantly higher methylation (median 78.3% vs. 50.9%, $p < 4.7 \times 10^{-182}$).

(D) Venn diagram illustrating the overlap between DMR-associated genes and DEGs identified between Disc-sFGR and Disc-N twins.

(E) Representative lollipop plots of CpG methylation levels and RNA-seq expression profiles at the *ATP7A* and *LIG4* loci in Disc-sFGR (blue) and Disc-N twins (red). Yellow shaded regions indicate DMRs located within promoter-proximal regions.

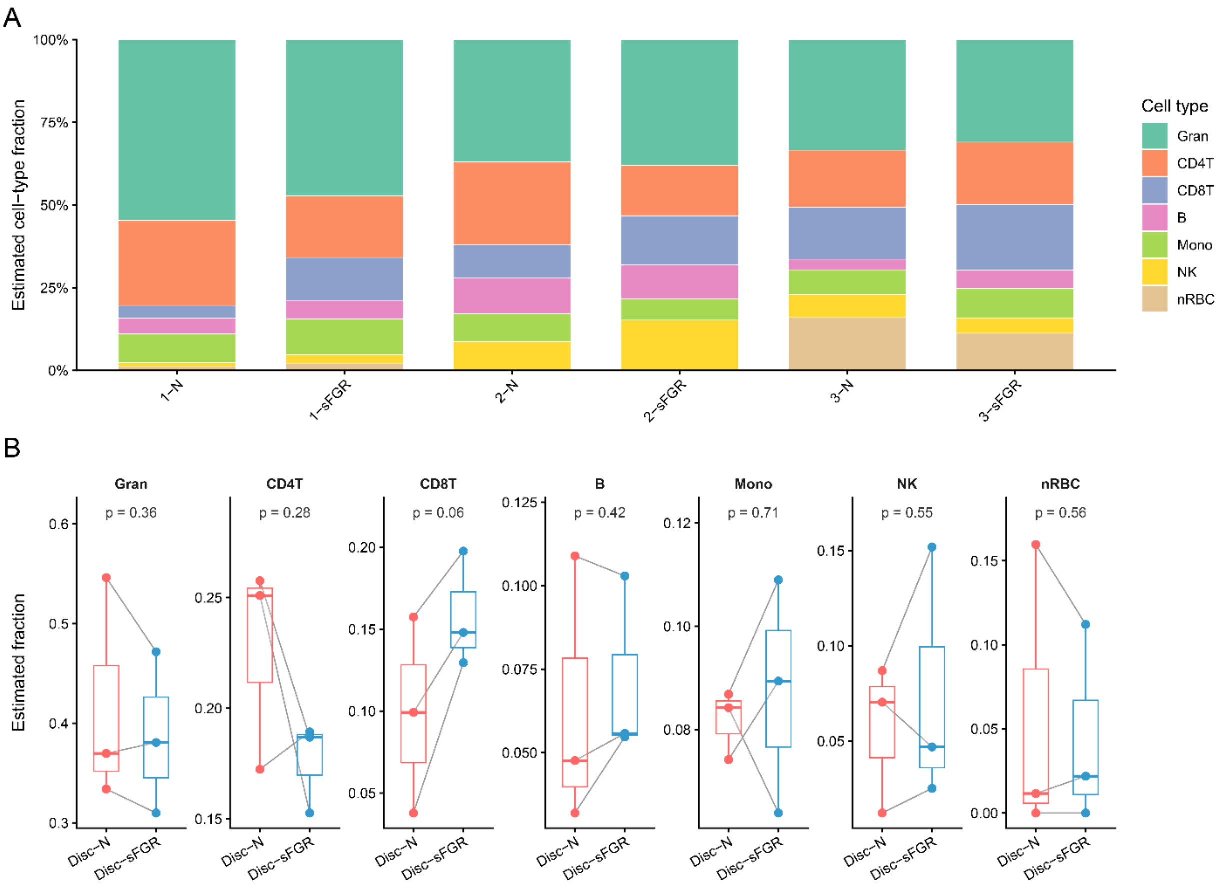

A
Estimated cell−type fraction
100%
75%
50%
25%
0%
1−N
1−sFGR
2−N
2−sFGR
3−N
3−sFGR
Cell type
Gran
CD4T
CD8T
B
Mono
NK
nRBC
B
Estimated fraction
Gran
p = 0.36
CD4T
p = 0.28
CD8T
p = 0.06
B
p = 0.42
Mono
p = 0.71
NK
p = 0.55
nRBC
p = 0.56
Disc−N
Disc−sFGR

**Figure S3. EpiDISH cell-type deconvolution of WGBS data in discordant twin pairs**

(A) Estimated cell-type fraction per sample using EpiDISH (Gran: granulocyte, CD4T: CD4+ T cell, CD8T: CD8+ T cell, B: B cell, Mono: monocyte, NK: natural killer cell, nRBC: nucleated red blood cell).

(B) Paired comparison of estimated cell-type fraction between Disc-N and Disc-sFGR twins. Grey lines connect twin pairs. P-values are from paired two-sided t-tests.
